# Survey of peridomestic mammal susceptibility to SARS-CoV-2 infection

**DOI:** 10.1101/2021.01.21.427629

**Authors:** Angela M. Bosco-Lauth, J. Jeffrey Root, Stephanie M. Porter, Audrey E. Walker, Lauren Guilbert, Daphne Hawvermale, Aimee Pepper, Rachel M. Maison, Airn E. Hartwig, Paul Gordy, Helle Bielefeldt-Ohmann, Richard A. Bowen

**Affiliations:** Colorado State University, Fort Collins, Colorado, USA; U.S. Department of Agriculture, Animal and Plant Health Inspection Service, Wildlife Services, National Wildlife Research Center, Fort Collins, Colorado, USA; School of Chemistry & Molecular Biosciences and Australian Infectious Diseases Research Centre, The University of Queensland. St Lucia, Queensland, Australia

**Keywords:** Coronavirus, cottontail rabbit, deer mouse, experimental infection, fox squirrel, ground squirrel, house mouse, peridomestic, mesocarnivore, raccoon, rodent, SARS-CoV-2, striped skunk, wildlife

## Abstract

Wild animals have been implicated as the origin of SARS-CoV-2, but it is largely unknown how the virus affects most wildlife species and if wildlife could ultimately serve as a reservoir for maintaining the virus outside the human population. Here we show that several common peridomestic species, including deer mice, bushy-tailed woodrats, and striped skunks, are susceptible to infection and can shed the virus in respiratory secretions. In contrast, we demonstrate that cottontail rabbits, fox squirrels, Wyoming ground squirrels, black-tailed prairie dogs, house mice, and racoons are not susceptible to SARS-CoV-2 infection. Our work expands upon the existing knowledge base of susceptible species and provides evidence that human-wildlife interactions could result in continued transmission of SARS-CoV-2.

## Introduction

The rapid global expansion of severe acute respiratory syndrome (SARS) coronavirus 2 (SARS-CoV-2) has been unprecedented in modern history. While the original human infection(s) were potentially linked to wild animals in a wet market (*1*), human-to-human transmission is currently the dominant mechanism of viral spread. Peridomestic animals, which are represented by wild and feral animals living within close proximity to humans, represent key species to evaluate for SARS-CoV-2 epidemiology for multiple reasons. First, given their common associations with humans and anthropogenically modified habitats, they represent the wildlife species with the greatest chance of exposure to the virus from humans (i.e., reverse zoonosis) or pets such as cats. Second, should select peridomestic wildlife prove to be susceptible to the virus and have the capacity to replicate it to high viral titers, these species would have the potential to maintain the virus among conspecifics. Third, should some species possess the maintenance host criteria mentioned above, they would represent wildlife species that would have the greatest chance (e.g., shedding ability and proximity to humans) to spread the virus back to humans. Wild rodents, cottontail rabbits (*Sylvilagus* sp.), raccoons (*Procyon lotor*) and striped skunks (*Mephitis mephitis*) can exhibit peridomestic tendencies in urban and suburban environments. Members of all these species/taxonomic groups have been shown to shed influenza A viruses following experimental inoculations (*2,3,4)*, suggesting they might harbor productive infections when exposed to other human-pathogenic respiratory viruses.

Based upon protein analyses of amino acid residues of ACE2, TMPRSS2 and S protein, species susceptibility analyses suggested that, among other taxonomic groups, both carnivores and wild rodents are potentially high-risk groups (*5,6,7).* Predicting specific species’ susceptibility, however, is more challenging. Looking at protein sequence analysis of ACE2 binding with the S protein of SARS-CoV-2, one study indicated that raccoons could be ruled out as potential hosts for SARS-CoV-2 (*6*) and a different study based upon sequence analysis suggested that the western spotted skunks (*Spilogale gracilis*) had a very low prediction of SARS-CoV-2 S-binding propensity (*7*). Similarly, the same study also suggested that American mink (*Neovison viso*n) have a similar prediction as western spotted skunks (*7)*. However, over the last several months, outbreaks of SAR-CoV-2 in commercial mink farms have been noted in Europe and more recently in the U.S. (*8,9*). Respiratory problems, rapid transmission, and/or unusually high mortality have been noted in this species in various regions (*9,10*), which suggests that the aforementioned analyses have limitations.

Rodents are the largest and most diverse order of mammals, so it is unsurprising that the susceptibility of rodents to SARS-CoV-2 varies by species. To date, only a handful of rodent species have been evaluated as potential reservoir hosts or animal models for SARS-CoV-2, and the results largely indicate that outbred species, including lab animals, are at most only moderately affected. Most non-transgenic laboratory mice (*Mus musculus*) are resistant to infection, while transgenic humanized mice and hamsters, including Syrian hamsters (*Mesocricetus auratus)* and dwarf hamsters (*Phodopus* sp.), are highly susceptible, with at least one report of Roborovky’s dwarf hamsters becoming fatally diseased within three days of exposure (*11,12,13).* Other species, including deer mice (*Peromyscus maniculatus),* become infected and shed low titers of virus, but the infection is subclinical (*14,15*). Considering that there are more than 1700 species of rodents world-wide, many of which exist closely at the human-wildlife interface, there remain many unanswered questions about SARS-CoV-2 and wild rodents.

Various lagomorphs exist as pets, livestock, and peridomestic wildlife, and as such are in prime position to come into contact with SARS-CoV-2 infected humans. In one study, New Zealand white rabbits were experimentally infected and shed infectious virus for up to seven days without signs of clinical disease (*16*) Wild rabbits, particularly cottontails in the U.S., are prolific and commonly found around human dwellings, farms, and commercial buildings. Further, as with rodents, wild rabbits are highly likely to be predated upon by domestic cats. Thus, determining the susceptibility of these animals is critically important to interpreting the risk posed to them and by them from infection with SARS-CoV-2.

Among carnivores, felids and mustelids have been frequently linked to SARS-CoV-2 infections since the early stages of the pandemic. Domestic cats are highly susceptible to SARS-CoV-2 and are capable of transmitting the virus to other cats, suggesting that they could potentially transmit to other animals as well (*17,18*). While striped skunks are currently considered to be mephitids, they are highly related to mammals within the family mustelidae and were formerly classified as mustelids. Thus, based on the findings of SARS-CoV-2 susceptibility in various mustelids, the closely related mephitids are a logical candidate to evaluate for the replication of this virus. Raccoons are notoriously associated with human environments and frequently interact with human trash and sewage, which has been proposed as a potential indirect means for infected humans to transmit SARS-CoV-2 to mammalian wildlife (e.g., raccoons and select mustelids) (*19,20,21*). Thus, it is important to determine the relative susceptibility of these common peridomestic carnivores and assess the likelihood that they could propagate infection.

In this study, we assessed six common peridomestic rodent species for susceptibility to SARS-CoV-2: deer mice, wild-caught house mice (*Mus musculus)*, bushy-tailed woodrats (aka “pack rats”; *Neotoma cinerea*), fox squirrels (*Sciurus niger)*, Wyoming ground squirrels (*Urocitellus elegans*), and black-tailed prairie dogs (*Cynomys ludovicianus*). These rodents are common in many parts of the United States, several of them frequently come into close contact with humans and human dwellings, and some are highly social animals, thus increasing the likelihood of pathogen transmission among conspecifics. In addition, we evaluated three other common peridomestic mammals: cottontail rabbits, raccoons, and striped skunks. Our results indicate that 33% (3/9) of the species evaluated are susceptible to SARS-CoV-2 infection, suggesting that wildlife may become critically implicated in the continued persistence of the virus.

## Materials and Methods

### Animals

The following mixed-sex animals were evaluated for susceptibility to SARS-CoV-2: Deer mice, house mice, bushy-tailed woodrats, Wyoming ground squirrels, black-tailed prairie dogs, fox squirrels, cottontail rabbits, striped skunks, and raccoons. Deer mice, house mice and bushy-tailed woodrats were trapped using Sherman traps baited with grain. Wyoming ground squirrels, fox squirrels, black-tailed prairie dogs, and cottontails were trapped using Tomahawk live traps (e.g., 7 x 7 x 20 or 7 x 7 x 24). All trapping was done in Northern Colorado (Larimer, Jackson and Weld counties) in accordance with Colorado wildlife regulations and with appropriate permits in place. Skunks and raccoons were purchased from a private vendor. Animals were housed in an Animal Biosafety Level-3 (ABSL3) facility at Colorado State University, in 12’x18’ rooms with natural lighting and controlled climate. Mice, black-tailed prairie dogs, and Wyoming ground squirrels were group housed by species with *ad libitum* access to water and food. All other animals were housed individually with access to food and water *ad libitum*. Rodents were maintained on Teklad^®^ Rodent Diet (Enviro, Madison, WI) supplemented with fresh fruit and occasional nuts. Rabbits were fed alfalfa pellets (Manna Pro^®^ Corp, Denver, Colorado) supplemented with grass hay and apples. Skunks and raccoons were maintained on Mazuri^®^ Omnivore Diet (Mazuri Exotic Animal Nutrition, St. Louis, MO) supplemented with fresh fruit and occasional eggs. Raccoons, striped skunks and black-tailed prairie dogs were implanted with thermally-sensitive microchips (Bio-Thermo Lifechips, Destron-Fearing) for identification and temperature measurement, deer mice were ear notched; all other animals were identified by cage number or distinct markings.

### Virus

SARS-CoV-2 virus strain WA1/2020WY96 was obtained from BEI Resources (Manassas, VA, USA), passaged twice in Vero E6 cells and stocks frozen at −80°C in Dulbecco’s Modified Eagle Medium (DMEM) with 5% fetal bovine serum and antibiotics. Virus stock was titrated on Vero cells using standard double overlay plaque assay (*17*) and plaques were counted 72 hours later to determine plaque-forming units (pfu) per ml.

### Virus challenge

Prior to challenge with SARS-CoV-2, most animals were lightly anesthetized as needed with 1-3 mg/kg xylazine and 10-30 mg/kg ketamine hydrochloride (Zetamine™) and a blood sample collected just before inoculation (Day 0). Virus diluted in phosphate buffered saline (PBS) was administered to all species via pipette into the nares (50ul for deer and house mice, 100ul for bushy-tailed woodrats, and 200ul for all other species) and animals were observed until fully recovered from anesthesia. Virus back-titration was performed on Vero cells immediately following inoculation, confirming that animals received between 4.5 and 4.9 log_10_ pfu of SARS-CoV-2.

### Sampling

Groups of three animals from each species (two for ground squirrels) were used for preliminary studies to evaluate viral shedding and acute pathological changes. For these animals, oral swabs were obtained pre-challenge and on days 1-3 post-challenge, at which time animals were euthanized and the following tissues harvested for virus isolation and formalin fixation: trachea, nasal turbinates, lung, heart, liver, spleen, kidney, small intestine, and olfactory bulb. The exception to this was raccoons, for which only one animal was euthanized at day 3; the remaining two were kept through day 28 to evaluate serological response. The remaining 3-6 animals per select species were swabbed daily from days 0-5 and 7 to further evaluate duration of viral shedding (if any). Striped skunks and raccoons were sedated for all sampling and a nasal swab was collected in addition to the oral swab. Tissues harvested from animals euthanized on day 7 were evaluated as for the day 3 animals. The remaining animals were euthanized at 28 days post-infection (DPI) and tissues were harvested for histopathology and serum was collected for serology. Table 1 illustrates the necropsy scheme for each species.

**Table 1.** Wildlife species evaluated for experimental infections with SARS-CoV-2 and day post infection the animals were euthanized.

| Animals | # euthanized at 3<br>DPI* | # euthanized at 7 DPI | # euthanized at 28<br>DPI |
| --- | --- | --- | --- |
| Deer mice (n=9) | 3 | 3 | 3 |
| House mice (n=6) | 3 | 0 | 3 |
| Bushy-tailed<br>woodrats(n=6) | 3 | 0 | 3 |
| Fox squirrels (n=3) | 3 | 0 | 0 |
| Wyoming ground<br>squirrels (n=2) | 2 | 0 | 0 |
| Black-tailed prairie dogs (n=9) | 3 | 3 | 3 |
| Cottontails (n=3) | 3 | 0 | 0 |
| Raccoons (n=3) | 1 | 0 | 2 |
| Striped skunks (n=6) | 3 | 0 | 3 |

\*Table footnotes: \*DPI = Days post-infection

### Clinical observations

Clinical evaluations were performed for all animals daily and included assessment for temperament and presence of any clinical signs of disease, such as ocular discharge, nasal discharge, ptyalism, coughing/sneezing, dyspnea, diarrhea, lethargy, anorexia, and if moribund. The stress of handling wild animals for sampling precluded the ability to obtain accurate body temperature measurements; as such, temperature was excluded in these preliminary studies for all species except skunks and raccoons, which were implanted with thermal microchips and could be measured under sedation during sampling.

### Viral assays

Virus isolation was performed on all oral swab, nasal swab and 3 DPI tissue samples by double overlay plaque assay on Vero cells as previously described (*17*). Briefly, 6-well plates with confluent monolayers of cells were washed once with PBS and inoculated with 100 μl of serial 10-fold dilutions of samples, incubated for 1 hour at 37°C, and overlaid with a 0.5% agarose in MEM containing 2% fetal bovine serum and antibiotics/antifungal agents. A second overlay with neutral red dye was added at 48 hours and plaques were counted at 72 hours. Viral titers were reported as the log_10_ pfu per swab (oropharyngeal/nasal) or per gram (tissue).

### Serology

Plaque reduction neutralization assays (PRNT) were performed as previously described (*17*). Serum was heat-inactivated for 30 mins at 56°C, and two-fold dilutions prepared in BA-1 (Tris-buffered MEM containing 1% bovine serum albumin) starting at a 1:5 dilution were aliquoted onto 96-well plates. An equal volume of virus was added to the serum dilutions and incubated for 1 hour at 37°C. Following incubation, serum-virus mixtures were plated onto Vero monolayers as described for virus isolation assays. Antibody titers were recorded as the reciprocal of the highest dilution in which >90% of virus was neutralized.

### qRT-PCR

Plaques were picked from culture plates from each positive animal to confirm SARS-CoV-2 viral shedding. RNA extractions were performed per the manufacturer’s instructions using Qiagen QiaAmp Viral RNA mini kits. RT-PCR was performed as recommended using the E_Sarbeco primer probe sequence as described by Corman and colleagues (*22*) and the Superscript III Platinum One-Step qRT-PCR system (Invitrogen), with the following modification: the initial reverse transcription was at 50°C. RNA standards for PCR were obtained from BEI Resources (Manassas, VA, USA).

### Histopathology

Animal tissues were fixed in 10% neutral-buffered formalin for 12 days and transferred to 70% ethanol prior to processing for paraffin-embedding, sectioning for H&E staining. Slides were read by a veterinary pathologist blinded to the treatments.

## Results

### Viral shedding

Of the nine species evaluated, three (deer mice, bushy-tailed woodrats, and striped skunks) shed infectious virus following challenge (Figure 1). Deer mice, which have previously been demonstrated to shed infectious SARS-CoV-2 experimentally (*15* Griffin), shed virus orally for up to four days and virus was isolated from lungs (n=3/3) and trachea (n=2/3) from animals harvested at 3 DPI. All nine inoculated deer mice shed virus on at least two of the first four days following infection, with peak titers of 3.1 log_10_ pfu/swab. Bushy-tailed woodrats shed virus orally for up to five days post inoculation (n=6/6) and virus was isolated from turbinates (n=2/3), trachea (n=1/3) and lung (n=1/3) from animals necropsied on 3 DPI. Peak titers from bushy-tailed woodrats reached 3.0 log_10_ pfu/swab by 3 DPI. Interestingly, the single bushy-tailed woodrat for which infectious virus was isolated from the lungs only shed 1.3 log_10_ pfu/swab orally on the day of necropsy, but the lungs contained 5.2 log_10_ pfu/gram virus. Striped skunks, which had to be handled under heavy sedation, were sampled on days 1-3, 5, and 7, during which time three of the six infected animals shed orally, nasally, or both, with one animal shedding up to 7 DPI. Of the three skunks necropsied on 3 DPI, two had infectious virus in the turbinates, but not in other tissues tested. One of those two animals had 3.2 log_10_ pfu/gram in the turbinates but failed to shed detectable virus nasally or orally prior to euthanasia. In general, viral titers were slightly higher in nasal samples compared to oral, but overall peak titers in skunks were relatively low, with oral titers reaching 2 log_10_ pfu/swab and nasal flush titers at 2.3 log_10_ pfu/swab. All animals with plaque-assay positive samples were confirmed for SARS-CoV-2 by RT-PCR. Similarly, all animals that were negative on plaque assay were confirmed negative for viral shedding by RT-PCR.

**Figure 1:**
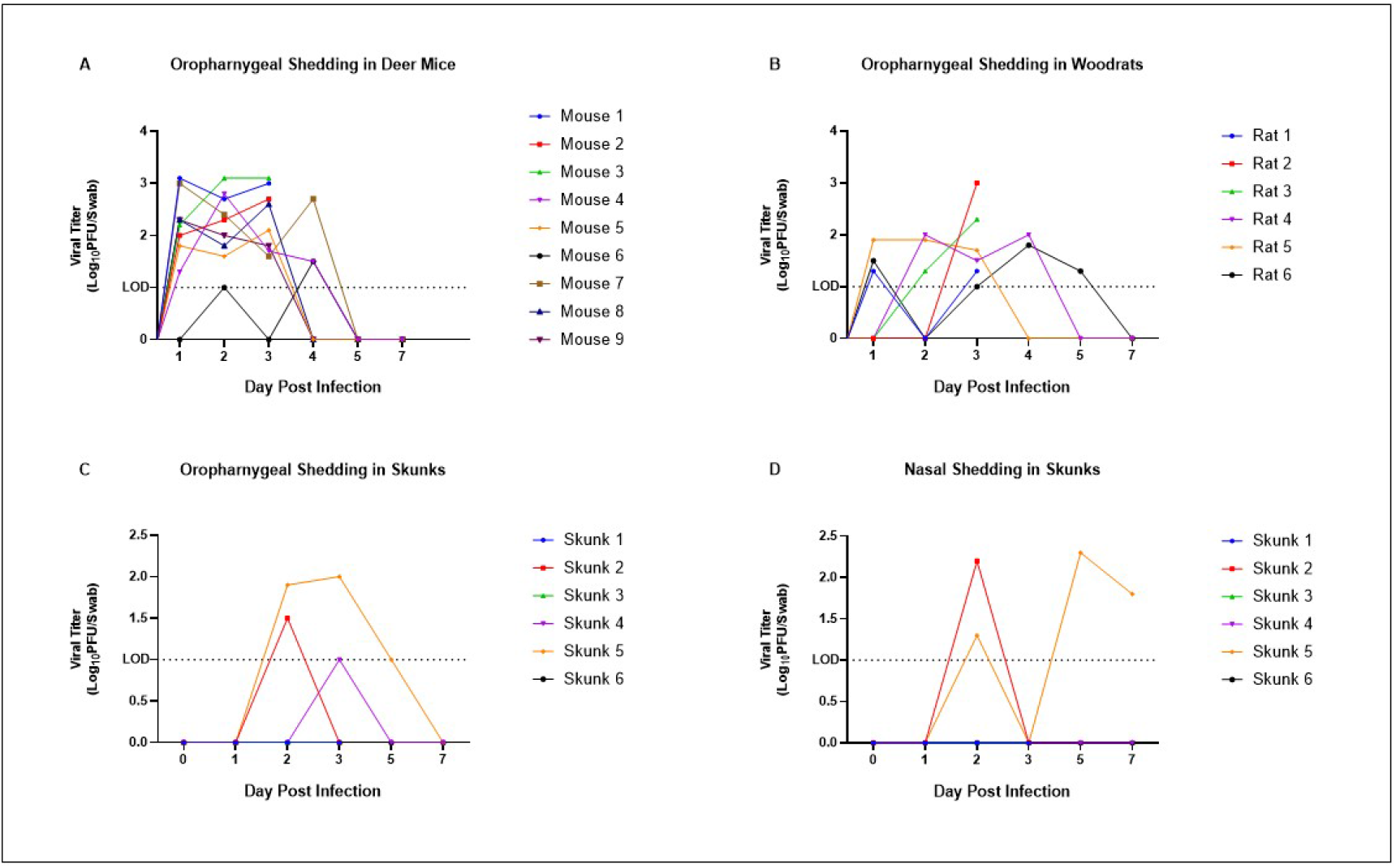
Oropharyngeal shedding of SARS-CoV-2 in deer mice (A), bushy-tailed woodrats (B) and striped skunks (C) and nasal shedding in striped skunks (D). Values expressed as log_10_ pfu/swab; limit of detection 1 log_10_ pfu.

### Seroconversion

All animals were seronegative against SARS-CoV-2 at the time of inoculation (<50% viral neutralization at 1:10 serum dilution). Based on the lack of evidence of infection and the overall difficulty of maintaining wildlife, we opted not to hold subsets of squirrels or rabbits for additional time to assess seroconversion. Neutralizing antibody titers were assessed in all animals euthanized at 28 DPI, which included deer mice, house mice, bushy-tailed woodrats, black-tailed prairie dogs, raccoons and striped skunks. All species which had detectable viral infections (deer mice, skunks, and bushy-tailed woodrats) also developed neutralizing antibodies, while the other species (house mice, raccoons, and black-tailed prairie dogs) did not. Deer mice and bushy-tailed woodrats reached or exceeded titers of 1:80, and the two skunks that shed infectious virus reached or exceeded titers of 1:160, while the single skunk that did not shed virus had a titer of 1:10 at 28 DPI. Animals euthanized at 3 DPI were not tested for seroconversion as previous investigations have demonstrated that neutralizing antibodies are typically not detectable during acute infection (*23*).

### Clinical disease

None of the animals exhibited clinical signs of disease (see methods for symptoms) at any time during the study. Skunks and raccoons, which were sedated for procedures which involved sampling, failed to display elevated temperatures at those times. In addition to clinical signs, behavior was monitored by observing animals through double-paned glass and assessing eating and response to provided enrichment (playing with toys, eating treats, using hides, etc.), and none of the animals were observed to behave abnormally following infection when compared to the acclimation period.

### Pathology

None of the animals had gross lesions at the time of necropsy. On histopathologic examination, rare, small foci of mild macrophage and neutrophil infiltration were noted in the lungs of two woodrats and two deer mice with one of the latter also having mild vasculitis. Two skunks presented with well-developed bronchiole associated lymphoid tissue (BALT), but inflammation was not apparent in the lungs or other tissues.

## Conclusions

COVID19 has had a significant impact on the human population globally, but so far very little is known about how SARS-CoV-2 virus impacts wildlife. Domestic cats and dogs have repeatedly been shown to be infected by SARS-CoV-2, but with few exceptions are asymptomatic or develop mild clinical disease (*17,24,25).* Farmed mink, on the other hand, are not only susceptible to infection, but can develop fulminating fatal disease (*10,26*). In contrast, ferrets, which are closely related to mink, shed virus following infection but the infection is subclinical (*27*). Raccoon dogs, which were heavily implicated in the SARS-CoV-1 outbreak in 2004, are susceptible to SARS-CoV-2 infection, but remain subclinical (*28*). Experimentally, deer mice can be infected and shed the virus via oral secretions, as demonstrated by this study and others (*14,15*). However, other mice, including wild house mice and non-transgenic laboratory strains of this species, are not susceptible to infection by SARS-CoV-2 (*29*). Studies in which bats and select small mammals were experimentally exposed to SARS-CoV-2 show that some species (i.e., fruit bats [*Rousettus aegyptiacus]* and tree shrews [*Tupaia belangeri])*, are capable of minimal viral replication while others (big brown bats [*Eptesicus fuscus])* do not appear to become infected at all, which suggests that while the virus may have originated in bats, they are unlikely to serve as reservoir hosts (*30,31,32*). The confounding clinical response to infection between closely related species makes predicting impacts on wildlife and their potential for reservoir maintenance difficult. Despite best attempts to predict host susceptibility based on receptor similarity or other modeling approaches, experimental infections remain the gold standard for evaluating the susceptibility of an animal to infection and following the course of disease.

Our results demonstrate that several common peridomestic wildlife species, including deer mice, bushy-tailed woodrats, and striped skunks are susceptible to SARS-CoV-2 infection and can shed infectious virus. Importantly, our work and the work of others indicate that so far, the majority of exposed wildlife species develop mild to no clinical disease and either fail to shed virus at all or shed low levels for short durations. Perhaps equally important is that these experimental infections suggest that we can rule out several common rodents, select wild lagomorphs and raccoons as potential SARS-CoV-2 reservoirs. There are, however, limitations to these experimental models, namely that the animals in our studies were directly exposed to high doses (e.g., 5 log_10_ pfu) of virus, which is unlikely to be representative of an exposure in nature. Additionally, experimental infections using low numbers of apparently healthy, immunocompetent animals do not generate sufficient data to fully characterize the risk posed to animals of varying ages and health status. However, the results of this work and the work of others, combined with the dramatic response to infection seen in certain species such as mink, indicate that the possibility exists of SARS-CoV-2 infecting wildlife, establishing a transmission cycle, and becoming endemic in non-human species. In particular, the relatively high titers observed in select woodrat tissues (e.g., 5.2 log_10_ pfu/gram of lung) suggests that a predator-prey transmission scenario among this rodent species and various small wild and domestic carnivore species is plausible. The major outcomes of such an event include direct threat to the health of wildlife and establishment of a reservoir host, which could complicate control measures of this virus in human populations. Experimental studies to identify and characterize species’ response to SARS-CoV-2 infection help scientists classify those species that are at highest risk and allow for the implementation of prevention measures. For example, both deer mice and bushy-tailed woodrats are commonly found in barns and sheds in very close proximity to humans, so when cleaning out sheds or attempting to rodent-proof barns, people should consider wearing appropriate personal protective equipment, both to prevent exposure to the pathogens rodents carry as well as to prevent exposing wildlife to SARS-CoV-2. Likewise, humans with COVID19 who also own cats and dogs should practice extra precaution with their pets, including minimizing the pet’s exposure to wildlife. Notably, a photo-monitoring study provided evidence that striped skunks can commonly use the same urban cover types (e.g., outbuildings and decks) as domestic cats (*33*). Intentionally available pet food and spilled bird feed, which were two of the attractants evaluated, produced instances where skunks and domestic cats were documented to be on study sites simultaneously or nearly simultaneously, which could facilitate interspecies transmission of SARS-CoV-2.

Wildlife and SARS-CoV-2 are intricately involved, from the initial spillover event to potential reverse zoonosis, and we will undoubtedly continue to discover more susceptible species as the search for zoonotic reservoirs continues. COVID19 is just the latest in a series of examples of how the human-wildlife interface continues to drive the emergence of infectious disease. The use of experimental research, surveillance, and modeling as tools for predicting outbreaks and epidemics will hopefully provide the knowledge base and resources necessary to prevent future pandemics.

## Acknowledgments

This work was funded by internal funding from Colorado State University and the U.S. Department of Agriculture, Animal and Plant Health Inspection Service. A. Walker and L. Guilbert were supported by the USDA Animal Health and Disease Veterinary Summer Scholar’s Program.

## Disclaimers

None

## Author Bio

Angela Bosco-Lauth is an Assistant Professor in the Department of Biomedical Sciences at Colorado State University. Here research interests include pathogenesis, transmission and ecology of infectious disease.

## References

1. Zhou P, Yang X-L, Wang X-G, Hu B, Zhang L, Zhang W, et al. A pneumonia outbreak associated with a new coronavirus of probable bat origin. Nature. 2020 Mar 1;579(7798):270–3.

2. Root JJ, Bosco-Lauth AM, Bielefeldt-Ohmann H, Bowen RA. Experimental infection of peridomestic mammals with emergent H7N9 (A/Anhui/1/2013) influenza A virus: Implications for biosecurity and wet markets. Virology. 2016 Jan;487:242–8.

3. Shriner SA, VanDalen KK, Mooers NL, Ellis JW, Sullivan HJ, Root JJ, et al. Low-Pathogenic Avian Influenza Viruses in Wild House Mice. Davis T, editor. PLoS ONE. 2012 Jun 15;7(6):e39206.

4. Romero Tejeda A, Aiello R, Salomoni A, et al. Susceptibility to and transmission of H5N1 and H7N1 highly pathogenic avian influenza viruses in bank voles (*Myodes glareolus*). Vet Res 46, 51 (2015).

5. Martínez-Hernández F, Isaak-Delgado AB, Alfonso-Toledo JA, Muñoz-García CI, Villalobos G, Aréchiga-Ceballos N, et al. Assessing the SARS-CoV-2 threat to wildlife: Potential risk to a broad range of mammals. Perspectives in Ecology and Conservation. 2020 Oct;18(4):223–34.

6. Luan J, Lu Y, Jin X, Zhang L. Spike protein recognition of mammalian ACE2 predicts the host range and an optimized ACE2 for SARS-CoV-2 infection. Biochemical and Biophysical Research Communications. 2020 May;526(1):165–9.

7. Damas J, Hughes GM, Keough KC, Painter CA, Persky NS, Corbo M, et al. Broad host range of SARS-CoV-2 predicted by comparative and structural analysis of ACE2 in vertebrates. Proc Natl Acad Sci USA. 2020 Sep 8;117(36):22311–22.

8. Oreshkova N, Molenaar RJ, Vreman S, Harders F, Oude Munnink BB, Hakze-van der Honing RW, et al. SARS-CoV-2 infection in farmed minks, the Netherlands, April and May 2020. Eurosurveillance [Internet]. 2020 Jun 11 [cited 2020 Dec 23];25(23).

9. DeLiberto T and Shriner S. Coronavirus disease 2019 update (536): Animal, USA (Utah), wild mink, first case. ProMED-mail, International Society for Infectious Diseases. Posted December 13, 2020.

10. Molenaar RJ, Vreman S, Hakze-van der Honing RW, Zwart R, de Rond J, Weesendorp E, et al. Clinical and Pathological Findings in SARS-CoV-2 Disease Outbreaks in Farmed Mink (*Neovison vison*). Vet Pathol. 2020 Sep;57(5):653–7.

11. Younes S, Younes N, Shurrab F, Nasrallah GK. Severe acute respiratory syndrome coronavirus-2 natural animal reservoirs and experimental models: systematic review. Rev Med Virol [Internet]. 2020 Nov 18 [cited 2021 Jan 18]; Available from: https://onlinelibrary.wiley.com/doi/10.1002/rmv.2196

12. Sia SF, Yan LM, Chin AWH, Fung K, Choy KT, Wong AYL, et al. Pathogenesis and transmission of SARS-CoV-2 in golden hamsters. Nature. 2020 Jul;583(7818):834–838.

13. Trimpert J, Vladimirova D, Dietert K, Abdelgawad A, Kunec D, Dökel S, et al. The Roborovski dwarf hamster is a highly susceptible model for a rapid and fatal course of SARS-CoV-2 infection. Cell Rep. 2020 Dec 8;33(10):108488.

14. Fagre A, Lewis J, Eckley M, Zhan S, Rocha SM, Sexton NR, et al. SARS-CoV-2 infection, neuropathogenesis and transmission among deer mice: Implications for reverse zoonosis to New World rodents [Internet]. Microbiology; 2020 Aug [cited 2020 Dec 7]. Available from: http://biorxiv.org/lookup/doi/10.1101/2020.08.07.241810

15. Griffin BD, Chan M, Tailor N, Mendoza EJ, Leung A, Warner BM, et al. North American deer mice are susceptible to SARS-CoV-2 [Internet]. Microbiology; 2020 Jul [cited 2020 Dec 14]. Available from: http://biorxiv.org/lookup/doi/10.1101/2020.07.25.221291

16. Mykytyn AZ, Lamers MM, Okba NMA, Breugem TI, Schipper D, van den Doel PB, et al. Susceptibility of rabbits to SARS-CoV-2. Emerging Microbes & Infections. 2021 Jan 10(1): 1–7.

17. Bosco-Lauth AM, Hartwig AE, Porter SM, Gordy PW, Nehring M, Byas AD, et al. Experimental infection of domestic dogs and cats with SARS-CoV-2: Pathogenesis, transmission, and response to reexposure in cats. Proc Natl Acad Sci USA. 2020 Oct 20;117(42):26382–8.

18. Shi J, Wen Z, Zhong G, Yang H, Wang C, Huang B, et al. Susceptibility of ferrets, cats, dogs, and other domesticated animals to SARS–coronavirus 2. Science. 2020 May 29;368(6494):1016–20.

19. Gross J, Elvinger F, Hungerford LL, Gehrt SD. Raccoon use of the urban matrix in the Baltimore Metropolitan Area, Maryland. Urban Ecosyst. 2012 Sep;15(3):667–82.

20. Hoffmann CO, Gottschang JL. Numbers, Distribution, and Movements of a Raccoon Population in a Suburban Residential Community. Journal of Mammalogy. 1977 Nov 29;58(4):623–36.

21. Franklin AB, Bevins SN. Spillover of SARS-CoV-2 into novel wild hosts in North America: A conceptual model for perpetuation of the pathogen. Science of The Total Environment. 2020 Sep;733:139358.

22. Corman VM, Landt O, Kaiser M, Molenkamp R, Meijer A, Chu DK, et al. Detection of 2019 novel coronavirus (2019-nCoV) by real-time RT-PCR. Eurosurveillance [Internet]. 2020 Jan 23 [cited 2020 Nov 28];25(3).

23. Kellam P, Barclay W. The dynamics of humoral immune responses following SARS-CoV-2 infection and the potential for reinfection. Journal of General Virology. 2020 Aug 1;101(8):791–7.

24. Patterson EI, Elia G, Grassi A, Giordano A, Desario C, Medardo M, et al. Evidence of exposure to SARS-CoV-2 in cats and dogs from households in Italy. Nat Commun. 2020 Dec;11(1):6231.

25. de Morais HA, dos Santos AP, do Nascimento NC, Kmetiuk LB, Barbosa DS, Brandão PE, et al. Natural Infection by SARS-CoV-2 in Companion Animals: A Review of Case Reports and Current Evidence of Their Role in the Epidemiology of COVID-19. Front Vet Sci. 2020 Oct 27;7:591216.

26. Hammer AS, Quaade ML, Rasmussen TB, Fonager J, Rasmussen M, Mundbjerg K, et al. SARS-CoV-2 Transmission between Mink (*Neovison vison*) and Humans, Denmark. Emerg Infect Dis [Internet]. 2021 Feb [cited 2020 Dec 7];27(2).

27. Kim Y-I, Kim S-G, Kim S-M, Kim E-H, Park S-J, Yu K-M, et al. Infection and Rapid Transmission of SARS-CoV-2 in Ferrets. Cell Host & Microbe. 2020 May;27(5):704–709.e2.

28. Freuling CM, Breithaupt A, Müller T, Sehl J, Balkema-Buschmann A, Rissmann M, et al. Susceptibility of Raccoon Dogs for Experimental SARS-CoV-2 Infection. Emerg Infect Dis. 2020 Dec;26(12):2982–5.

29. Cohen J. From mice to monkeys, animals studied for coronavirus answers. Science. 2020 Apr 17;368(6488):221–2.

30. Hall JS, Knowles S, Nashold SW, Ip HS, Leon AE, Rocke T, et al. Experimental challenge of a North American bat species, big brown bat (*Eptesicus fuscus*), with SARS-CoV-2. Transboundary and Emerging Diseases. 2020 Dec 9;tbed.13949.

31. Schlottau K, Rissmann M, Graaf A, Schön J, Sehl J, Wylezich C, et al. SARS-CoV-2 in fruit bats, ferrets, pigs, and chickens: an experimental transmission study. The Lancet Microbe. 2020 Sep;1(5):e218–25.

32. Zhao Y, Wang J, Kuang D, Xu J, Yang M, Ma C, et al. Susceptibility of tree shrew to SARS-CoV-2 infection. Sci Rep. 2020 Dec;10(1):16007.

33. Weissinger MD, Theimer TC, Bergman DL, Deliberto TJ. Nightly and seasonal movements, seasonal home range, and focal location photo-monitoring of urban striped skunks (Mephitis mephitis): Implications for rabies transmission. Journal of Wildlife Diseases. 2009 Apr;45(2):388–97.

